# Selective enrichment of high-affinity clade II N_2_O-reducers in a mixed culture

**DOI:** 10.1101/2024.02.09.579283

**Authors:** Michele Laureni, Francesc Corbera Rubio, DaeHyun Daniel Kim, Savanna Browne, Nina Roothans, David G. Weissbrodt, Karel Olavaria, Nadieh de Jonge, Sukhwan Yoon, Martin Pabst, Mark C.M. van Loosdrecht

## Abstract

Microorganisms encoding for the N_2_O reductase (NosZ) are the only known biological sink of the potent greenhouse gas N_2_O, and are central to global N_2_O mitigation efforts. Yet, the ecological constraints selecting for different N_2_O-reducers strains and controlling the assembly of N_2_O-respiring communities remain largely unknown. Of particular biotechnological interest are clade II NosZ populations, which usually feature high N_2_O affinities and often lack other denitrification genes. Two planktonic N_2_O-respiring mixed cultures were enriched under limiting and excess dissolved N_2_O availability to assess the impact of substrate affinity and N_2_O cytotoxicity, respectively. Genome-resolved metaproteomics was used to infer the metabolism of the enriched populations. We show that clade II N_2_O-reducers outcompete clade I affiliates for N_2_O at sufficiently low sludge dilution rates (0.006 h^-1^), a scenario previously only theorized based on pure-cultures. Under N_2_O limitation, all enriched N_2_O-reducers encoded and expressed only clade II NosZ, while also possessing other denitrification genes. Two *Azonexus* and *Thauera* genera affiliates dominated the culture. We explain their coexistence with the genome-inferred metabolic exchange of cobalamin intermediates. Conversely, under excess N_2_O, clade I and II populations coexisted. Notably, the single dominant N_2_O-reducer (genus *Azonexus*) expressed most cobalamin biosynthesis marker genes, likely to contrast the continuous cobalamin inactivation by dissolved cytotoxic N_2_O concentrations (400 µM). Ultimately, we demonstrate that the solids dilution rate controls the selection among NosZ clades, albeit the conditions selecting for genomes possessing the sole *nosZ* remain elusive. Additionally, we suggest the significance of N_2_O-cobalamin interactions in shaping the composition of N_2_O-respiring microbiomes.

## Introduction

Nitrous oxide (N_2_O) is a potent greenhouse gas, with a global warming potential almost 300 times higher than CO_2_, and it is the predominant ozone-depleting substance in the atmosphere (1). Globally, N_2_O emissions have been increasing at a rate of 17 Tg N y^-1^ over the last decade (2). The rise is expected to continue if no mitigation efforts are put in place (1). Most of the produced N_2_O results from microbially mediated reactions in managed and engineered ecosystems characterized by high nitrogen loads, such as agricultural soils and wastewater treatment plants (WWTP) (3, 4). Consequently, robust emission mitigation strategies strongly rely on our understanding of the microbiology underlying N_2_O production and consumption.

N_2_O is produced primarily as a by-product of the first nitrification step, *i*.*e*. the biological oxidation of ammonia (NH_3_) to nitrite (NO_2_^-^), or during incomplete denitrification (5, 6). Denitrification is the sequential reduction of dissolved nitrate (NO_3_^-^) and NO_2_^-^ to dinitrogen gas (N_2_), with nitric oxide (NO) and N_2_O as obligate free intermediates (7). Each reduction step is catalysed by distinct enzymes and can function as an independent energy-conserving reaction (4, 7, 8). Denitrification can be carried out by a single microorganism (termed “generalist”) or a consortium of cooperating “specialists”, each of them performing one or few nitrogen oxides reductions (9). As such, denitrification can act both as a source or a sink of N_2_O depending on the genetic and metabolic repertoire of the microbial community members, and the environmental conditions (10, 11). Importantly, the last step of the denitrification pathway, the reduction of N_2_O to N_2_ catalysed by the N_2_O-reductase (NosZ), is the only known biological N_2_O sink (4).

Taxonomically, the ability to reduce N_2_O is widely distributed among diverse microbial groups (12), and the *nosZ* gene evolved in two phylogenetically distinct lineages, namely clade I and II (4). Interestingly, clade II NosZ appears to be more often encoded in genomes of non-denitrifying (specialist) N_2_O-reducers, *i*.*e*. microorganisms lacking other denitrifying enzymes and thus representing possible net N_2_O sinks (4, 13, 14). Limited pure culture-based observations suggests that clade II organisms feature higher affinities for N_2_O than clade I populations, albeit at usually lower maximum specific growth rates (15, 16). On these grounds, irrespective of their denitrifying capacity, clade II N_2_O-reducers hold the potential to further curb emissions by reducing residual dissolved N_2_O concentrations.

Denitrifying communities dominated by N_2_O-reducing specialists (17) and rich in clade II affiliates are ubiquitous (9, 12, 18-23). The abundance of clade II populations has been reported to inversely correlate with N_2_O emissions from soils (10, 24, 25). High relative abundances of clade II N_2_O-reducers have been obtained in lab-grown biofilms (15, 26, 27). A clear positive correlation between clade II *nosZ* gene abundance and longer solids retention times has been reported in the seminal chemostat work by Conthe and co-workers (28, 29). Yet, to date no enrichment has resulted in the selection of sole clade II organisms, and different traits such as the preferred electron donor (30), the sensitivity to other electron acceptors like NO_3_^-^ (31) and O_2_ (30), or the required presence of other nitrogen oxides (32) may also contribute to clades selection. Resolving the ecological constraints controlling the selection of different N_2_O-reducers would greatly benefit both biotechnological designs and our understanding of the biosphere response to a rapidly changing climate.

We postulate that the high solids dilution rates used in previous studies prevented the selection of clade II N_2_O-reducers (28, 29). To verify our hypothesis, two N_2_O-respiring mixed cultures were enriched under limitation of either electron donor (acetate) and electron acceptor (N_2_O), and an ultrafiltration membrane was used to impose a solids dilution rate five times lower than ever tested. The taxonomy and metabolic potential of the dominant community members at steady-state were identified via genome-resolved metagenomics. Shotgun metaproteomics was used to estimate the relative biomass contribution and infer the actual metabolism of each organism. Under N_2_O-limiting conditions we successfully cultivated a mixed culture where all N_2_O-reducers encoded and expressed the sole clade II NosZ, while still the conditions were not selective enough for specialist N_2_O-reducers. Additionally, metagenomic and metaproteomic data suggest a potential role of vitamin B_12_ cross-feeding in the assembly of and niche differentiation within N_2_O converting communities, which has not been previously reported.

## Materials and Methods

### Continuous enrichments

Two identical glass, continuous-flow membrane bioreactors (MBR) with a working volume of 2 L (Applikon, Delft, the Netherlands) were operated with acetate and N_2_O as sole electron donor and acceptor, respectively. To allow for direct comparisons, the set-up was identical to the one used by Conthe and co-workers (29). The sole difference was the use of a custom-made, submerged ultrafiltration membrane module (33) to uncouple the solids retention time (SRT) and the hydraulic retention time (HRT). In both MBRs, the hydraulic retention time (HRT) was maintained at 2.8 ± 0.3 d. The SRT was set at 6.9 ± 1.0 d, equivalent to a biomass dilution rate (D) of 0.006 ± 0.001 h^-1^ (Table 1). The temperature was controlled at 20 ± 1 °C, and the pH was maintained at 7.0 ± 0.1 with 1M HCl and 1M NaOH. Stirring at 750 rpm was ensured by two six-bladed flat turbines. Two independent 5850S mass-flow controllers (Brooks, Philadelphia, PA) were used in each MBR to mix pure N_2_O and N_2_, and achieve the target overall gas flow and N_2_O concentration (Figure 1). The carbon and nitrogen sources were provided separately. The two mineral media contained per litre: 90.5 mmol NaCH_3_COO·3H_2_O or 26.4 mmol NH_4_Cl, respectively, additionally to 7.4 mmol KH_2_PO_4_, 2.1 mmol MgSO_4_·7H_2_O, 0.5 mmol NaOH, 2 mg yeast extract and 2.5 ml of trace element solution (34). The first MBR operated under acetate limitation and N_2_O excess (hereafter referred to as *N*_*2*_*Oexc*) was inoculated directly with activated sludge from the Harnaschpolder wastewater treatment plant (Delft, the Netherlands). Excess N_2_O conditions, defined as no detectable acetate in the effluent, were achieved with influent acetate and N_2_O loads of 31.6 ± 2.0 and 382 ± 25 mmol/d, respectively, after a 30-days start-up phase. The second MBR operated under N_2_O limitation (*N*_*2*_*Olim*) was inoculated with biomass collected from *N*_*2*_*Oexc* after 98 days of operation. N_2_O limiting conditions in *N*_*2*_*Olim* were reached on day 60 and maintained until day 95 at influent acetate and N_2_O loads of 46.1 ± 1.5 and 84 ± 2.1 mmol/d, respectively (Figure 1). Occasional growth on the MBR walls and membrane was cleaned weekly.

**Table 1.** Biomass, ammonium, and N_2_O stoichiometric yields of the *N*_*2*_*Oexc* and *N*_*2*_*Olim* enrichments. Values were calculated based on the measured consumption and production rates after data reconciliation (days 100-150 for *N*_*2*_*Oexc* and 60-100 for *N*_*2*_*Olim*; Table S1), and standard deviations were estimated by propagating the relative error of each measured rate. The yields obtained by (29) at higher dilutions with an identical set-up, used for the estimation of the biomass-specific substrate consumption rate for maintenance, are also presented.

| D | N<sub>2</sub>O/Ac (inf.) | Y<sub>X/Ac</sub> | Y<sub>X/N<sub>2</sub>O</sub> | Y<sub>N<sub>2</sub>O/Ac</sub> | Y<sub>NH<sub>4</sub>/X</sub> |  |
| --- | --- | --- | --- | --- | --- | --- |
| h<sup>-1</sup> | mol/mol | mol/mol<sub>C</sub> | mol/mol | mol/mol | mol/mol |  |
| $0.006 \pm 0.001$ | $12.8 \pm 1.5$ | $0.15 \pm 0.03$ | $0.09 \pm 0.02$ | $3.42 \pm 0.35$ | $0.41 \pm 0.15$ | **N<sub>2</sub>Oexc** |
| $0.028 \pm 0.001$ | 15.9 | $0.25 \pm 0.02$ | $0.16 \pm 0.01$ | $3.06 \pm 0.2$ | $0.28 \pm 0.02$ | Conthe et al. 2018c |
| $0.089 \pm 0.003$ | 26.1 | $0.27 \pm 0.01$ | $0.17 \pm 0.01$ | $3.1 \pm 0.17$ | $0.28 \pm 0.01$ | Conthe et al. 2018c |
| $0.006 \pm 0.001$ | $1.8 \pm 0.1$ | $0.24 \pm 0.04$ | $0.15 \pm 0.03$ | $3.1 \pm 0.15$ | $0.44 \pm 0.13$ | **N<sub>2</sub>Olim** |
| $0.027 \pm 0.001$ | 2.3 | $0.32 \pm 0.04$ | $0.22 \pm 0.04$ | $2.86 \pm 0.51$ | $0.26 \pm 0.01$ | Conthe et al. 2018c |
| $0.086 \pm 0.003$ | 2.7 | $0.33 \pm 0.03$ | $0.26 \pm 0.02$ | $2.49 \pm 0.29$ | $0.25 \pm 0.02$ | Conthe et al. 2018c |

**Figure 1.**
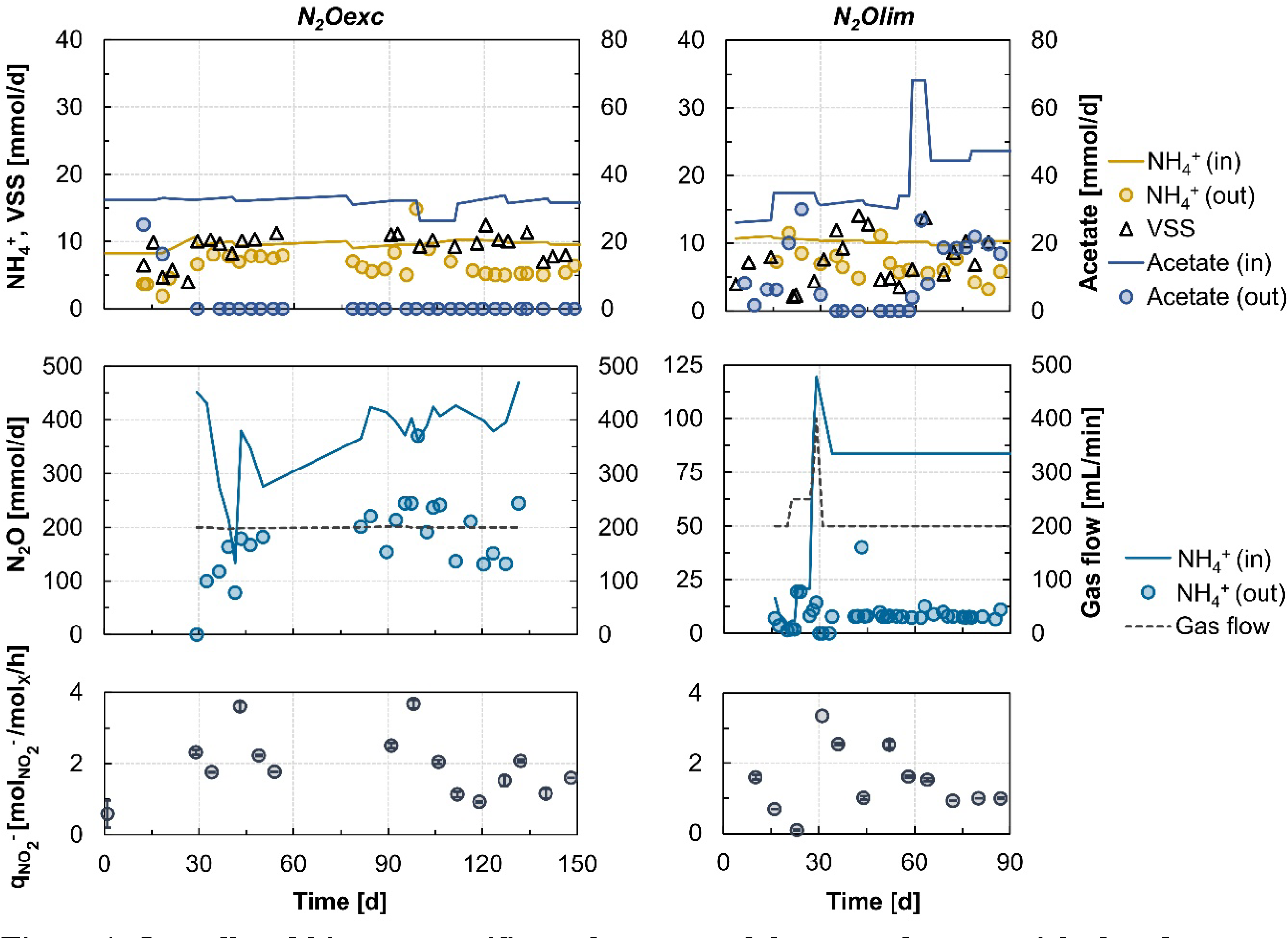
Overall and biomass specific performance of the two cultures enriched under excess (*N*_*2*_*Oexc*) and limiting (*N*_*2*_*Olim*) availability of nitrous oxide as sole electron donor. Influent and effluent acetate and ammonium flow rates, and biomass production rate. *Red triangle*: day of biomass sampling for metagenomic and metaproteomic analysis (**a, e**). Influent and effluent N_2_O rates, and imposed gas flow (**b, f**). Ex-situ maximum biomass specific NO_2_^-^ consumption rates (**c, g**).

### Denitrification batch tests

We aimed to quantify the long-term dynamics of the denitrification potential of the enrichments respiring exclusively N_2_O. To this end, the maximum biomass-specific nitrite reduction rate (q_NO2-_; mmol_NO2-_·mmol_X_^-1^·h^-1^) was used as denitrification proxy and quantified in batch. Fresh reactor effluent was collected under anoxic conditions, via continuous N_2_-sparging of the effluent collection vessel with N_2_ gas, and centrifuged at 4200 rpm for 20 min at room temperature. The recovered pellet was re-suspended in N_2_-sparged base mineral medium without ammonium or acetate, with the same composition as detailed above, to a biomass concentration of 0.9 ± 0.2 g_VSS_·L^-1^. Subsequently, 50 mL of cell suspension were aliquoted in 112 mL serum bottles. The bottles were sealed with rubber stoppers, and anoxic conditions were achieved by flushing with N_2_ for 20 min. After overnight incubation, appropriate volumes of anoxic stock-solutions of NH_4_Cl, NaCH_3_COO·3H_2_O and NaNO_2_ were added to reach the target initial concentrations of 2.1 mmol NH_4_^+^·L^-1^, 1.2 mmol CH_3_COO^-^ L^-1^, and 1.1 mmol NO_2_^-^·L^-1^. The frequency of bulk liquid sampling (4 mL) depended on biomass activity. Samples were immediately centrifuged for 5 minutes (4200 rpm, 4 °C), and the supernatant was kept for further analysis. The volumetric NO_2_^-^ consumption rate was calculated by linear regression of at least four data points, and average and standard deviations were obtained from duplicates. The biomass-specific reduction rates were calculated by dividing the volumetric rates by the biomass concentration, and standard deviations were calculated from the individual values and standard deviations using linear error propagation. Initial and final acetate concentrations were measured to confirm non-limiting conditions. All tests were performed in an Incubator Hood TH30 (Edmund Bühler GmbH, Bodelshausen, Germany) at 20 °C and 150 rpm. Negative controls with autoclaved biomass were also included. The pH remained in the range 7-8 without external control.

### Analytical procedures and calculations

Samples for influent and effluent supernatant analysis were centrifuged at 4ºC and 13000 rpm for 5 min, stored at -20 ºC, and analysed within less than 6 days. NH_4_^+^, NO_2_^-^ and NO_3_^-^ concentrations were measured spectrophotometrically with the Gallery™ Discrete Analyzer (Thermo Fisher Scientific). Acetate concentration was measured by high-performance liquid chromatography (Vanquish Core HPLC, Thermo Fisher Scientific) using an Aminex HPX-87H column (300 × 7.8 mm) (Bio-Rad, Waters Chromatography B.V.), calibrated with standard solutions ranging from 0 to 250 mM. In- and off-gas N_2_O concentrations were continuously monitored online by a Rosemount NGA 2000 off-gas analyser (Emerson). Before reaching the analyser, the off-gas was dried in a condenser operated with water at 4 °C using a cryostat bath (Lauda). N_2_O concentrations in the in- and off-gas of *N*_*2*_*Oexc* exceeded the off-gas analyser range, thus weekly grab-samples were measured with a CP-3800 gas chromatograph (Varian). The concentrations of total and volatile suspended solids (VSS, TSS) in the mixed liquors were determined according to standard methods (35), and an average biomass composition of CH_1.8_O_0.5_N_0.2_ (36) was assumed for downstream molar calculations. The stoichiometric yields (Table 1) were calculated from the steady-state volumetric and biomass specific conversion rates (Table S1) and the reactor operation measurements (Figure 1) after data reconciliation using the software Macrobal (37). Dissolved N_2_O concentrations were estimated from the average off-gas N_2_O concentrations with a Henry’s constant of 2.4·10^-4^ mmol/(L·Pa) (38).

### Metagenomics

<u>*Metagenome sequencing*</u>. Samples for metagenomic analysis were taken on days 123 (*N*_*2*_*Oexc*) and 83 (*N*_*2*_*Olim*). DNA was extracted using the DNeasy UltraClean Microbial Kit (Qiagen, The Netherlands). Library preparation and metagenomic sequencing were performed by Novogene Ltd. (Hongkong, China) on HiSeq platform (Illumina Inc., CA). Libraries were generated from 1 μg DNA per sample using the NEBNext Ultra DNA Library Prep Kit (NEB #E7645, USA) following the manufacturer’s instructions. DNA was fragmented by sonication to a size of 350 bp, and DNA fragments were end-polished, A-tailed, and ligated with a full-length adapter for further PCR amplification. PCR products were purified (AMPure XP system), and libraries were quantified using real-time PCR after size distribution analysis using an Agilent 2100 Bioanalyzer. Index-coded samples were clustered with a cBot Cluster Generation System according to the manufacturer’s instructions. Ultimately, libraries were sequenced to generate 150 bp paired-end reads (HiSeq sequencing platform, Illumina Inc., US). <u>*Metagenome assembly and binning*</u>. The raw shotgun metagenome reads were screened using Trimmomatic v0.36 with the parameters set as follows: *LEADING:3 TRAILING:3 SLIDINGWINDOW:4:15 MINLEN:70* (39). The trimmed reads were assembled *de novo* into contigs using metaSPAdes v3.14.0 with default parameters (40). The trimmed reads were mapped back onto the assembled contigs using the *mem* algorithm implemented in BWA v0.7.17, and the mapping data were sorted using SAMtools v1.10 (41, 42). The coverage of each contig was calculated using *jgi_summarize_bam_contig_depths* command implemented in MetaBat2 with the parameters *minContigLength* and *minContigDepth* set to 2000 and 2, respectively (43). These coverage data were used as the input to MetaBat2 for binning the contigs into metagenome assembled genomes (MAG) (43). CheckM v1.1.2 software was implemented to assess the quality of the bins by computing their completeness and contamination (*lineage_wf* command) (44). MAGs potentially belonging to a single organism were identified and subsequently merged (*merge* command) after manual inspection of their marker gene sets and coverage information (44). The bins were subsequently refined using refineM, removing the contigs violating consistency in terms of contig statistics (computed with *scaffold_stats* command) and/or taxonomic affiliations of the protein-coding sequences within (45). Protein-coding sequences were also predicted with refine using the *call_genes* command and their taxonomic affiliation was inferred using the *taxon_profile* command with the GTDB database release 80 as reference. Subsequently, inconsistent contigs were identified with the *taxon_filter* and removed. After filtering out all of potential outlier contigs, the bins were further filtered with the completeness and contamination thresholds set to 75% and 5%, respectively. <u>*Metagenome annotation*</u>. The taxonomy affiliations of the refined MAGs were inferred using GTDB-tk v2.3.2 with GTDB database release 214 (46). To estimate the relative abundances of the organisms represented by the MAGs, the quality-trimmed raw reads were mapped onto the contigs in the finalized MAGs using BWA MEM v0.7.17 with default parameters (41). The alignment files were sorted with SAMtools v1.10, and the coverages of the contigs in the MAGs were calculated using *coverage* command of checkM v1.1.2 (44). The relative abundance of each MAG was calculated using *profile* command and represented as the percentage of total reads mapped to the contigs in the MAG, normalized by the total size of the contigs in the MAG. The potential coding sequences within each MAG were annotated using MetaErg v1.2.0 with Pfam, TIGRFAMS, Metabolic-hmms, SwissProt, and RefSeq databases as reference (47). Annotation was supplemented with Ghostkoala (48). The functions, domain structures, KEGG Orthology (KO) numbers, Gene Orthology numbers (GO), and Enzyme Commission numbers (EC) were included in the annotation. The annotation of the two *nosZ* clades was manually refined on Interpro (49) based on the twin-arginine translocation (clade I, IPR006311) or the general secretory (clade II, IPR026468) pathway-specific signal peptides (50). The cobalamin-dependent epoxyqueuosine reductase *queG* (EC 1.17.99.6), and its cobalamin-independent functional homologue *queH* (51), were differentiated phylogenetically using annotated *queG* and *queH* sequences from UniProtKB (52). The same approach was used to differentiate the two nucleotide loop assembly marker genes for cobalamin production, *cobP, cobU* and *cobY*.

### Mass spectrometry based proteomics

<u>*Protein extraction and proteolytic digestion*</u>. *Protein Extraction:* Approximately 25 mg of biomass (wet weight) was collected in an Eppendorf tube and solubilized in a suspension buffer consisting of 175 µL B-PER reagent (Thermo Scientific) and 175 µL TEAB buffer (50 mM TEAB, 1% (w/w) NaDOC, adjusted to pH 8.0). Next, 0.1 g of glass beads (acid-washed, approximately 100 µm in diameter) were added and the cells were disrupted using 5 cycles of bead beating on a vortex for one minute, followed by cooling on ice for 30 seconds. Then, a freeze/thaw step was performed by freezing the suspension at -80°C for 20 minutes and thawing it while shaking in an incubator. The cell debris was pelleted at 14,000 x g for 10 minutes while being cooled at 4°C. The supernatant was transferred to a new Eppendorf tube. *Protein Precipitation*: The protein was precipitated by adding 1 volume of TCA (trichloroacetic acid) to 4 volumes of the supernatant. The solution was then incubated at 4°C for 10 minutes and pelleted again at 14,000 x g for 10 minutes. The resulting protein pellet was washed twice with 200 µL of ice-cold acetone. *Trypsin Digestion*: The protein pellet was dissolved in 200 mM ammonium bicarbonate containing 6M urea to achieve a final concentration of approximately 100 µg/µL. To 100 µL of the protein solution, 30 µL of a 10 mM DTT solution was added and incubated at 37°C for one hour. Next, 30 µL of a freshly prepared 20 mM IAA solution was added and incubated in the dark for 30 minutes. The solution was then diluted to a concentration of <1 M Urea using a 200 mM ammonium bicarbonate buffer, and an aliquot of approximately 25 µg of protein was digested with Trypsin (Promega, Trypsin to protein ratio of 1:50) at 37°C overnight. *Solid Phase Extraction*: The peptides were desalted using an Oasis HLB 96-well plate (Waters) as per the manufacturer’s instructions. The purified peptide eluate was then dried using a speed vac concentrator. <u>*Large-scale shotgun proteomics and label-free quantification.*</u> An aliquot containing approximately 250 ng protein digest was analysed in duplicates using an one-dimensional shotgun proteomics approach. Each duplicate was injected twice. Briefly, the samples were analysed using a nano-liquid-chromatography system consisting of an EASY nano-LC 1200, equipped with an Acclaim PepMap RSLC RP C18 separation column (50 µm × 150 mm, 2µm), and an QE plus Orbitrap mass spectrometer (Thermo Scientific, Germany). The flow rate was maintained at 350 nL/min. Following sample loading and a wash step at 5% B, a linear gradient was performed, first to 25% B over 88 minutes, and further to 50% B over additional 25 minutes. Finally, the separation column was back equilibration to starting conditions. Solvent A was H_2_O containing 0.1% formic acid, and solvent B consisted of 80% acetonitrile in H_2_O and 0.1% formic acid. The spectrometer was operated in the data dependent acquisition mode, thereby measuring peptide signals from the 385-1250 m/z range at 70K resolution with the AGC target of 3e6 and max IT of 100 ms. The top 10 signals were isolated at a window of 2.0 m/z and fragmented using a NCE of 28. Fragments were acquired at 17K resolution, at an AGC target of 5e4 and 100 ms max IT. <u>*Database searching, comparison of conditions and data visualisation*</u>. Data were analysed with a database of *in silico* translated gene-coding sequences obtained from shotgun metagenome experiments of the analysed enrichment cultures (as described above), using PEAKS Studio 10 (Bioinformatics Solutions Inc, Canada). Database searching was performed by allowing for 20 ppm parent ion mass error tolerance and 0.02 m/z fragment ion mass error tolerance and allowing of up to 2 missed cleavages. Carbamidomethylation was set as fixed modification and methionine oxidation and N/Q deamidation were considered as variable modifications. Peptide spectrum matches were filtered against 1% false discovery rate (FDR) and protein identifications with ≥ 2 unique peptides were considered as significant identifications. The relative abundance of each protein is presented in terms of normalized spectral counts, defined as spectral counts per protein divided by its molecular weight, and divided by the sum of spectral counts of the specific injection. Only proteins identified in at least two out of four injections were included in the analysis, and the average of their normalized spectral counts in the different injections is taken as proxy for their relative abundance. RStudio v2023.03.1 (53) with R v4.3.0 (54) was used for data analysis and visualization.

### Data availability

Raw DNA reads were deposited on the NCBI Sequence Read Archive and medium- and high-quality MAGs were deposited in Genbank under BioProject PRJNA1054980. The mass spectrometry proteomics raw data, reference sequence database and database search files have been deposited in the ProteomeXchange consortium database with the dataset identifier PXD030677.

## Results and Discussion

### Reduced biomass growth under continuous N_2_O excess

Two N_2_O-respiring mixed cultures were successfully enriched in a chemostat over multiple generations with acetate and N_2_O as sole electron donor and acceptor, respectively (Figure 1; Table S1). Continuous-flow membrane reactors were used to select for microbial communities based on substrate affinity under low-growth-rate conditions. The first community (*N*_*2*_*Oexc*) was enriched directly from activated sludge under N_2_O excess, defined as undetectable acetate in the reactor effluent. The second community (*N*_*2*_*Olim*) was inoculated with biomass collected from *N*_*2*_*Oexc* after 100 days of operation, and run under N_2_O limitation at a residual acetate concentration of 23.7 ± 5.5 mmol/L. The solids dilution rate (D) was set to 0.006 h^-1^, to verify our hypothesis that previous attempts to select for clade II organisms have been hindered by excessively high solids washouts. The imposed D was lower than the growth rate reported for clade II *nosZ*-possessing *Anaeromyxobacter dehalogenans* 2CP-C (0.0076 h^-1^), the slowest-growing N_2_O-reducing bacterium examined by (16), and the minimum dilution rate (0.027 h^-1^) that has been applied in previous non-axenic N_2_O-reducing chemostat works (29).

The biomass growth yields on N_2_O (Y_X/N2O_) and acetate (Y_X/Ac_) were lower than those obtained by (29) with an identical set-up and comparable operating conditions except for the use of an ultrafiltration membrane to operate at four times lower solids dilution rates (Table 1). The observed lower yields are consistent with higher substrate requirements for maintenance at lower growth rates. The stoichiometric consumption of 3.1-3.4 moles N_2_O per mole acetate consumed was slightly higher than the values obtained by (29) (Table 1). The yields of ammonium consumed per biomass produced (Y_NH4/X_) were higher than expected for the commonly assumed empirical biomass composition (CH_1.8_O_0.5_N_0.2_ (36)), yet comparable to the ones of the *Pseudomonas stutzeri* JM300 strain grown on N_2_O and acetate (29). Importantly, the growth yields on both N_2_O and acetate were substantially lower in *N*_*2*_*Oexc* than in *N*_*2*_*Olim* (Table 1). This is in agreement with (29). This is likely related to the potential cytotoxic effects of N_2_O (55-57). Based on average off-gas N_2_O concentrations, the estimated dissolved N_2_O concentration in *N*_*2*_*Oexc* was 382±25 µM, exceeding by far the concentration at which cytotoxic effects have been previously reported (57). The negative impact of N_2_O on growth is further supported by a 2.4-fold higher specific acetate consumption rate for maintenance, namely 0.019 ± 0.001 (*N2Oexc*) and 0.008 ± 0.001 (*N2Olim*) C-mol_Ac_·mol_X_^-1^·h^-1^, as estimated by linear regression of the growth yields on acetate over the D range covered by this study and (29) (Figure S1).

### *Rhodocyclaceae*-dominated N_2_O-respiring enrichments

The metagenomes of the two steady-state enrichments were sequenced to resolve the identity and functional potential of the enriched organisms. A total of 75.2·10^6^ and 64.5·10^6^ reads (*N*_*2*_*Oexc* and *N*_*2*_*Olim*, respectively) were obtained after quality filtering, and their assembly yielded 6550 and 7001 contigs (> 2000 bp) with N50 values of 18.8·10^3^ and 12.6·10^3^ bp. Contigs were binned, resulting in 10 and 17 metagenome-assembled genomes (MAGs), with completeness > 75% and contamination < 5

% (Table 2). The recovered MAGs accounted for 94.5 and 74.1 % of the quality-filtered reads in *N*_*2*_*Oexc* and *N*_*2*_*Olim*, respectively. The obtained binning coverage indicates that the recovered MAGs represent the majority of the two microbiomes. In terms of relative reads abundance, Bin.820_exc (*N*_*2*_*Oexc*) and Bin.9_lim (*N*_*2*_*Olim*) were the dominant MAGs, and accounted for about 83.2 and 67.1 % of the reads in the respective reactors. Low quality MAGs were grouped with the unbinned reads (Table 2).

**Table 2.** Genomic and proteomic characteristics of the recovered MAGs. MAGs marked with (*) affiliate with the *Rhodocyclaceae* family.

|  |  | Matagenome |  |  |  |  |  |  | Metaproteome |  |  |
| --- | --- | --- | --- | --- | --- | --- | --- | --- | --- | --- | --- |
| Bin ID |  | Comple. | Contam. | Bin size [Mbp] | N50 [bp] | GC [%] | Mapped reads [%] | Predicted proteins | Detected proteins | Rel. abund. spectra [%] | Taxonomy |
| N<sub>2</sub>O<sub>exc</sub> | Bin.820_exc* | 97.6 | 0.0 | 2.7 | 171993 | 59.8 | 83.2 | 2614 | 1077 | 75.8 | genus *Azonexus* |
|  | Bin.2_exc | 94.5 | 0.4 | 3.2 | 49531 | 60.5 | 2.0 | 3009 | 143 | 5.4 | genus *Phaeovulum* |
|  | Bin.24_exc | 97.5 | 0.5 | 3.3 | 33922 | 59.3 | 1.5 | 3023 | 117 | 5.1 | genus *Pseudorhodobacter* |
|  | Bin.15_exc | 98.2 | 0.0 | 3.5 | 133854 | 46.6 | 5.6 | 3299 | 98 | 3.7 | genus *Bdellovibrio* |
|  | Bin.3_exc | 77.8 | 1.8 | 3.1 | 20050 | 65.3 | 0.4 | 3014 | 20 | 0.8 | genus *Aquamicrobium* |
|  | Bin.12_exc | 98.9 | 0.1 | 3.1 | 74431 | 33.7 | 0.3 | 2787 | 6 | 0.3 | species *Flavobacterium filum* |
|  | Bin.18_exc | 99.5 | 0.0 | 2.7 | 81605 | 40.4 | 0.7 | 2424 | 8 | 0.3 | genus *Moheibacter* |
|  | Bin.16_exc | 99.0 | 0.0 | 2.5 | 57459 | 37.3 | 0.3 | 2262 | 8 | 0.2 | genus *Kaistella* |
|  | Bin.17_exc | 99.0 | 0.0 | 2.9 | 65957 | 39.2 | 0.3 | 2480 | 3 | 0.1 | genus *Ferruginibacter* |
|  | Bin.9_exc | 95.8 | 0.5 | 2.5 | 13365 | 35.0 | 0.2 | 2281 | - | 0.0 | genus *Ferruginibacter* |
|  | Unbinned | - | - | 26.4 | 5341 | - | 5.5 | 27394 | 133 | 8.4 | - |
| N<sub>2</sub>O<sub>lim</sub> | Bin.9_lim* | 87.3 | 0.0 | 2.7 | 32281 | 63.5 | 67.1 | 2492 | 648 | 50.8 | genus *Azonexus* |
|  | Bin.16_lim* | 100.0 | 0.7 | 4.2 | 38798 | 69.5 | 1.9 | 3747 | 181 | 13.5 | species *Thauera phenylacetica* |
|  | Bin.14_lim | 77.2 | 0.1 | 2.0 | 4921 | 40.5 | 0.1 | 2090 | 43 | 2.1 | family *Weeksellaceae* |
|  | Bin.21_lim | 99.5 | 0.3 | 2.6 | 104725 | 35.2 | 0.8 | 2170 | 60 | 1.8 | genus *Ferruginibacter* |
|  | Bin.6_lim | 75.4 | 1.8 | 2.5 | 24195 | 48.9 | 0.4 | 2366 | 17 | 0.7 | species *Seleniivibrio woodruffii* |
|  | Bin.24_lim | 78.0 | 1.1 | 1.3 | 57096 | 24.0 | 0.8 | 1137 | 12 | 0.5 | phylum *Patescibacteria* |
|  | Bin.20_lim* | 76.9 | 0.9 | 3.1 | 6309 | 66.0 | 0.2 | 3184 | 7 | 0.4 | species *Azovibrio restrictus* |
|  | Bin.26_lim | 89.8 | 3.9 | 2.4 | 13434 | 61.7 | 0.2 | 2332 | 1 | 0.2 | genus *Pseudoflavonifractor* |
|  | Bin.11_lim | 93.0 | 1.6 | 3.9 | 17980 | 43.9 | 0.3 | 3203 | 6 | 0.1 | order *Bacteroidales* |
|  | Bin.2_lim | 93.5 | 0.1 | 1.7 | 11915 | 50.4 | 0.4 | 1697 | 4 | 0.1 | family *Anaerovoracaceae* |
|  | Bin.8_lim | 78.5 | 2.8 | 2.6 | 4471 | 38.0 | 0.2 | 2495 | 3 | 0.1 | genus *Niabella* |
|  | Bin.10_lim | 93.7 | 1.1 | 2.9 | 42970 | 33.1 | 0.4 | 2266 | 3 | 0.1 | order *Bacteroidales* |
|  | Bin.1_lim | 76.2 | 0.1 | 1.1 | 4590 | 37.2 | 0.1 | 1208 | 1 | 0.1 | genus *Paracholeplasma* |
|  | Bin.27_lim | 81.2 | 1.9 | 3.4 | 6337 | 37.4 | 0.2 | 3430 | 1 | 0.0 | genus *Fusibacter_C* |
|  | Bin.23_lim | 96.2 | 1.1 | 2.4 | 106442 | 31.2 | 0.7 | 1972 | 3 | 0.0 | order *Bacteroidales* |
|  | Bin.22_lim | 79.2 | 1.1 | 1.3 | 5309 | 41.6 | 0.1 | 1457 | - | 0.0 | family *Erysipelotrichaceae* |
|  | Bin.18_lim | 97.3 | 0.0 | 2.4 | 37598 | 34.4 | 0.2 | 2000 | - | 0.0 | family *Paludibacteraceae* |
|  | unbinned | - | - | 14.9 | 5762 | - | 25.9 | 15000 | 334 | 29.5 | - |

The proteomes of the same samples were analysed to quantify the relative biomass contribution of each microorganism represented by a MAG, and infer its actual metabolic role within the microbiome (58). Per sample, we obtained an average of 20981 and 16832 MS2 spectra (*N*_*2*_*Oexc* and *N*_*2*_*Olim*, respectively) over the quadruplicate injections. We identified an average of 14490 and 12449 peptides, resulting in 1613 and 1324 proteins identified with a FDR < 0.01 by at least 2 unique peptides in at least 2 out of 4 injections. The relative abundance of a protein was calculated by dividing its normalized spectral counts by the sum of the normalized spectral counts in the corresponding injection. About 91.7 and 74.7 % of the proteins identified in *N*_*2*_*Oexc* and *N*_*2*_*Olim*, respectively, were mapped to the recovered MAGs, providing a comprehensive coverage of the expressed functional potential of the enrichments. Bin.820_exc and Bin.9_lim accounted for 70.1 and 50.6 % of their respective metaproteome, consistent with their metagenomic-based relative abundance (Table 2). In contrast, Bin.16_lim, that represented only 1.9 % of the metagenomic reads in *N*_*2*_*Olim*, accounted for 14.0 % of the metaproteome, suggesting a more prominent metabolic role than predicted based on the sole metagenome. Moreover, for the three dominant MAGs we identified 1077 (Bin.820_exc), 648 (Bin.9_lim), and 181 (Bin.16_lim) proteins representing the 41.2, 26.0, and 4.8 % of their respective metagenome-predicted coding genes.

Taxonomically, the recovered draft genomes were taxonomically affiliated to the phyla *Bacteroidota* (*Bacteroidetes*; 12), *Pseudomonadota* (*Proteobacteria*; 7), *Bacillota (Firmicutes;* 5), *Bdellovibrionota* (1), *Patescibacteria* (1), and *Deferribacterota* (1) (Table 2; Table S2). All three dominant MAGs (Bin.820_exc, Bin.9_lim, and Bin.16_lim) affiliated with the family *Rhodocyclaceae*, and shared an average nucleotide identity < 84 % indicating that they do not belong to the same species (59). Bin.820_exc and Bin.9_lim affiliated with the genus *Azonexus*, while Bin.16_lim with the genus *Thauera*. Globally, the microbially diverse and metabolically versatile *Rhodocyclaceae* family constitutes a core taxon in WWTP microbiomes, driving denitrification and likely N_2_O reduction across diverse ecosystems (9, 12, 60-62). *Rhodocyclaceae* members are also commonly co-enriched based on their N_2_O reducing capability under non-axenic conditions in chemostats (28, 29, 63, 64) and biofilm-based systems (15). Recently, *Azonexus* genus affiliates have been enriched and isolated from biogas digestate, and their catabolic preference for N_2_O reduction has been shown both at kinetic and proteomic level (65). Our study further supports the prominent role of *Rhodocyclaceae* family members in providing a potential N_2_O sink in diverse ecosystems.

### N_2_O limitation and low biomass dilution select for high-affinity clade II N_2_O-reducers

Consistently with nitrous oxide being the sole electron acceptor, the nitrous oxide reductase (NosZ) was among the ten most abundant proteins with an assigned KO number in *N*_*2*_*Oexc* and *N*_*2*_*Olim*, accounting for 1.4 and 1.1 % of the total normalized spectral counts respectively (Figure 2; Table S2). All recovered MAGs under N_2_O excess possessed a *nosZ* gene. Bin.820_exc harboured genes encoding for the clade II NosZ protein, and contributed the most to its overall expression (Figure 2). Clade I harbouring genomes were also present in *N*_*2*_*Oexc*, albeit at very low relative abundances (Figure 2). The co-existence of the two clades aligns with the majority of characterized natural communities (4, 66) and N_2_O-respiring enrichments (15, 26, 27, 29, 64), and suggests that the imposed conditions were not selective enough for an absolute dominance of clade II N_2_O-reducers. In contrast, N_2_O limitation resulted in the sole enrichment of clade II N_2_O-respiring microorganisms, as further confirmed by the absence of the clade I *nosZ* gene sequence even within the unbinned reads (Figure 2). The NosZ protein belonging to the most abundant Bin.9_lim and Bin.16_lim shared about 80% of sequence identity with the NosZ of Bin.820_exc (*N*_*2*_*Oexc*). To the best of our knowledge, this constitutes the first enrichment of sole clade II *nosZ* harbouring organisms in non-axenic mixed microbial communities.

**Figure 2.**
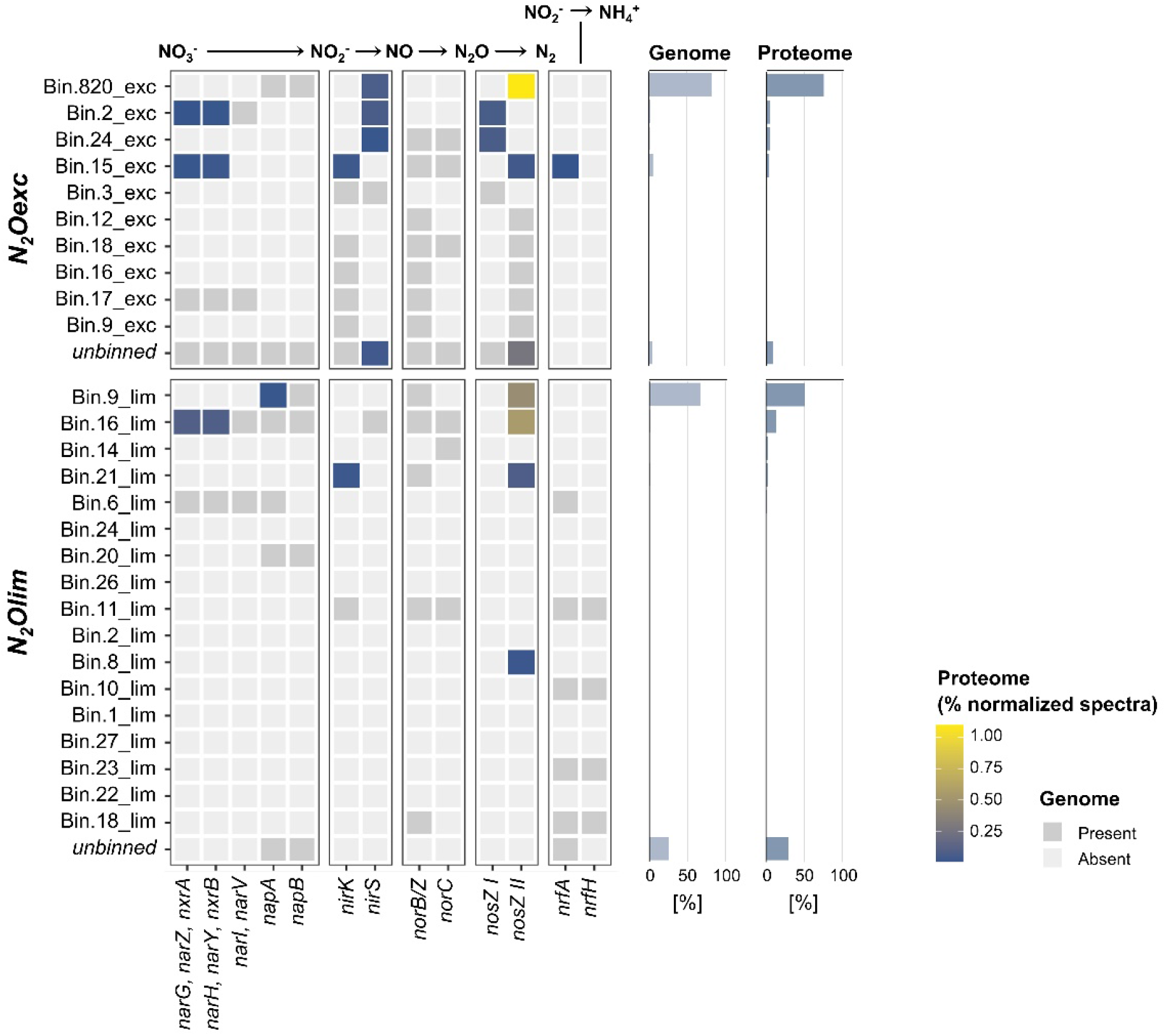
Denitrification and dissimilatory nitrite reduction to ammonium (DNRA). Gene presence (**dark grey tiles**) and protein abundance (**coloured tiles**) for the high- and medium-quality (completeness > 75 % and contamination < 5 %) metagenome assembled genomes (MAGs) enriched under excess (*N*_*2*_*Oexc*) and limiting (*N*_*2*_*Olim*) nitrous oxide availability. The sum of the normalized spectral counts for each protein was taken as a proxy for protein abundance. **Right bar charts**: relative abundance of each MAG in the metagenome (based on relative reads alignment) and the metaproteome (summed relative abundance of normalized spectral counts of peptides matching to predicted proteins in each MAG). **Abbreviations**: *napAB*: electron transfer (*napB*; EC:1.9.6.1; K02567) and catalytic (*napA*; K02568) subunits of the periplasmic nitrate reductase. *narGHI*: membrane-bound nitrate reductase, comprising the transmembrane protein *narI* (EC:1.7.5.1, 1.7.99.-; K00374) mediating the electron transfer from the quinol pool via *narH* (EC:1.7.5.1, 1.7.99.-; K00371) to the catalytic subunit *narG* (EC:1.7.5.1, 1.7.99.-; K00370). *nirK*: copper-containing nitrite reductase (EC:1.7.2.1; K00368). *nirS*: cytochrome *cd*_*1*_-containing nitrite reductase (EC:1.7.2.1, 1.7.99.1; K15864). *norCB*: nitric oxide reductase comprising the two subunits *norC* (K02305) and *norB/Z*; the latter comprises both the cytochrome c-dependent (*cNorB*; EC:1.7.2.5; K04561) and the menaquinol-dependent (*qNorB*/*NorZ*; EC:1.7.5.2) variants (68). *nrfAH*: catalytic (*nrfA*; EC:1.7.2.2; K03385) and electron donating (*nrfH*; K15876) subunits of the periplasmic dissimilatory cytochrome c nitrite reductase.

Contrary to our expectations, all *nosZ* encoding genomes were not specialist N_2_O-reducers and possessed also at least a nitrate or nitrite reductase gene, with the only exception of the low-abundant medium-quality (76.4 % completeness) Bin.8_lim (Figure 2). The periplasmic nitrate reductase (*napAB*) was encoded by all three dominant MAGs (Bin.820_exc, Bin.9_lim, Bin.16_lim), while only Bin.16_lim also possessed the membrane-bound dissimilatory nitrate reductase *narGHI*. In agreement with published denitrifiers genomes (9), MAGs possessing the cytochrome *cd*_*1*_-type nitrite reductase (*nirS*) did not encode the copper-containing nitrite reductase (*nirK*), and vice versa, with the only exception of the low-abundant Bin.3_exc (*N*_*2*_*Oexc*). Yet, none of the *nir* was annotated in Bin.9_lim (*N*_*2*_*Olim*). In contrast to often observed co-occurrence patterns (4, 9), no evident correlation emerged between the presence of *nirS* and *nosZ*, or the nitric oxide reductase (*cNorB* or *qNorB*). The ammonification potential, identified by the presence of the periplasmic energy-conserving periplasmic cytochrome c nitrite reductase *nrfAH* complex, was almost exclusively restricted to the N_2_O-limiting conditions with acetate excess (*N*_*2*_*Olim*), in an interesting parallel with the electron donor-rich environments selecting for ammonifiers (67). Despite N_2_O being the only electron acceptor, some NarG/NapA and NirK/NirS enzymes were detected in the proteome, yet at more than one order of magnitude lower abundances than NosZ (Figure 2). The observed constitutive expression of the denitrifying enzymes is consistent with the significant maximum *ex-situ* nitrite reduction capacity maintained in both enrichments (Figure 1) in analogy to (29) and (25).

Taken together, these results show that N_2_O limiting conditions at sufficiently low growth rates (here 0.006 h^-1^) exclusively select for *clade II* N_2_O-reducers. To date, this was only hypothesized based on the higher affinity displayed by a limited number of characterized clade II isolates (15, 16), and the positive correlation between clade II *nosZ* gene abundance and long retention times reported in the work of Conthe and colleagues (28, 29). Yet, the conditions further selecting for non-denitrifying, specialist N_2_O-reducers in natural and engineered microbiomes remain to be identified.

### Cobalamin auxotrophy as competitive advantage at high N_2_O

We postulated that the lower growth yields in *N*_*2*_*Oexc* resulted from the cytotoxic effects of elevated N_2_O levels in the growth medium. Consequently, this played an important role in the differences in microbial selection and community assembly of the two enrichments. N_2_O is known to selectively inactivate cobalamin (vitamin B_12_), an essential cofactor in bacterial metabolism (55, 56). High N_2_O concentrations have been shown to hinder microbial growth by compromising the activity of cobalamin-dependent enzymes (57). Cobalamin synthesis itself constitutes a significant genetic and metabolic burden for the cell, involving up to thirty enzymes (69), and in environmental communities it is usually performed by a small cohort of prototrophic low-abundant microorganisms (70, 71). The recovered MAGs were interrogated for genes encoding proteins involved in cobalamin synthesis and transport, and in core cobalamin-dependent metabolic pathways.

We identified Bin.820_exc (*N*_*2*_*Oexc*) and Bin.16_lim (*N*_*2*_*Olim*) as *de novo* cobalamin producers based on the presence of seven of the eight experimentally-verified biosynthetic marker genes proposed by (71) (corrin ring biosynthesis: *cbiL*/*cobI, cbiF*/*cobM, cbiC*/*cobH*; nucleotide loop assembly: *cobQ/cbiP, cobC1/cobC/cbiB/cobD, cobS/cobV* and *cobP/cobU*, except *cobY*; Figure 3, Table S3). The entire cobalamin production pathway was also almost completely annotated in both MAGs, and the majority of predicted proteins were detected in Bin.820_exc. In contrast, Bin.9_lim (*N*_*2*_*Olim*) contained all nucleotide loop assembly steps, but lacked two of the downstream corrin ring biosynthetic marker genes, namely *cbiL*/*cobI* and *cbiC*/*cobH*, along with a more inconsistent pathway annotation (Figure 3). On these grounds, while missing annotations *e*.*g*. due to non-homologous replacements cannot be ruled out, Bin.9_lim seems to have relied on the uptake of late cobalamin intermediates (*e*.*g*. cobinamide) either from the yeast extract present in the influent or produced by Bin.16_lim (71). The high protein expression of the outer membrane cobalamin/cobinamide transporter (BtuB; (69, 70)) in both MAGs further supports this hypothesis (Figure 3). Conversely, in *N*_*2*_*Oexc*, Bin.820_exc was devoid of the vitamin B12 transporter gene *btuB*, and the BtuB protein was detected at low abundances only in side populations.

**Figure 3.**
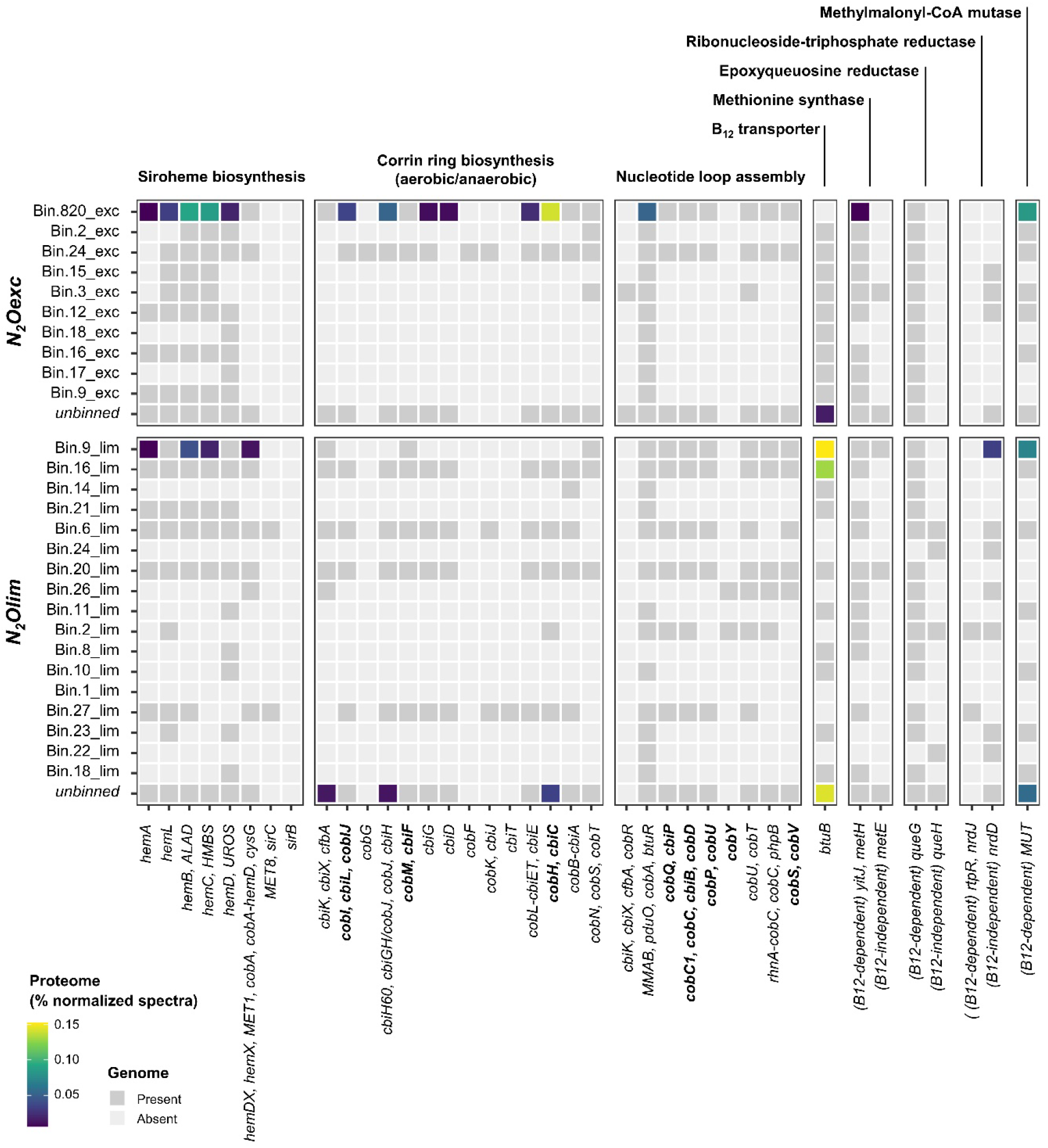
Cobalamin bio-synthesis and transport, and cobalamin (in)dependent enzymes. Gene presence (**dark grey tiles**) and protein abundance (**coloured tiles**) for the high- and medium-quality MAGs enriched under excess (*N*_*2*_*Oexc*) and limiting (*N*_*2*_*Olim*) nitrous oxide availability. **Annotations. Corrin ring biosynthesis (aerobic/anaerobic) marker genes:** *cobI, cbiL, cobIJ* (EC:2.1.1.130, EC:2.1.1.151; K03394, K13540). *cobM, cbiF* (EC:2.1.1.133, EC:2.1.1.271: K05936). cobH, cbiC (EC:5.4.99.61, EC:5.4.99.60; K06042). **Nucleotide loop assembly marker genes**: *cobQ, cbiP (EC:*6.3.5.10; K02232). *cobC1, cobC, cbiB, cobD* (EC:6.3.1.10; K02225, K02227). *cobP, cobU* (EC:2.7.1.156; K02231). *cobS, cobV* (EC:2.7.8.26; K02233). *cobY* (EC:2.7.7.62; K19712). **Transport and (in)dependent enzymes**. *btuB*: cobalamin/cobinamide transporter (K16092). *yitJ, metH*: cobalamin-dependent methionine synthase (EC:2.1.1.13, EC:1.5.1.54; K00548, K24042). *metE*: cobalamin-independent methionine synthase (EC:2.1.1.14; K00549). *queG*: cobalamin-dependent epoxyqueuosine reductase (EC:1.17.99.6; K18979). *queH*: cobalamin-independent epoxyqueuosine reductase (EC:1.17.99.6; K09765). *nrdJ, rtpR*: cobalamin-dependent ribonucleoside-triphosphate reductase (EC:1.17.4.2; K00524, K00527). *nrdD*: cobalamin-independent ribonucleoside-triphosphate reductase (EC:1.1.98.6; K21636). *MUT*: methylmalonyl CoA mutase (EC:5.4.99.2; K01847). Details for all bio-synthesis genes can be found in Table S3.

Both enrichments encoded the cobalamin-dependent processes most widely found in bacterial genomes, encompassing amino acid and nucleotide synthesis (70, 71). All three dominant MAGs possessed the cobalamin-dependent methionine synthase (*metH*), while Bin.9_lim encoded also its cobalamin-independent orthologue (*metE*) (Figure 3). *metE* has been previously shown to be up-regulated under cobalamin-limiting conditions, *e*.*g*. at high N_2_O, albeit displaying significantly lower activities (57). Yet, only the cobalamin-dependent MetH was detected in the proteome of Bin.9_lim. The reliance of all three MAGs on vitamin B12 was further supported by the presence of the cobalamin-dependent epoxyqueuosine reductase (*queG*), involved in tRNA modification, and by the expression of methylmalonyl CoA mutase (*MUT*), catalysing the synthesis of succinyl-CoA (72).

We posit that *de novo* cobalamin production provided Bin.820_exc with a selective advantage in *N*_*2*_*Oexc*. High N_2_O concentrations likely compromised both the internal and exogenous (*i*.*e*. from yeast extract) active vitamin B12 pool, making cobalamin self-sufficiency an essential metabolic trait. Conversely, the more permissive conditions in *N*_*2*_*Olim* made cobalamin cross-feeding viable, and its production dispensable (73). The high expression of the cobalamin transporter (BtuB) may suggest the reliance of both Bin.9_lim and Bin.16_lim on the uptake of the exogenous yeast extract cobalamin. However, we reason that the genome-inferred cobalamin-prototrophy provided Bin.9_lim with a higher relative fitness, *i*.*e*. allowing for an optimized cellular proteome allocation (74-76), while creating the ecological niche for the unexpected co-enrichment of the putative cobalamin-producer Bin.16_lim. Overall, the abundances of the corresponding proteins were relatively low especially in the *N*_*2*_*Olim* (Figure 3), in line with reported low detections of cobalamin production gene-transcripts even in cobalamin-free cultures (77). While caution is warranted in their interpretation, the higher detection of cobalamin synthesis proteins in *N*_*2*_*Oexc* compared to *N*_*2*_*Olim* (Figure 3) further emphasizes an increased need to sustain an otherwise continuously compromised cobalamin pool. Ultimately, cobalamin cross-feeding may underpin the often reported coexistence of N_2_O reducers in natural and engineered complex ecosystems.

## Acknowledgements

The authors are deeply indebted to Dimitry Sorokin (TU Delft) for the thorough discussions and invaluable insights. ML was supported by a Marie Skłodowska-Curie Individual Fellowship (grant agreement 752992) and a VENI grant from the Dutch Research Council (NWO) (project number VI.Veni.192.252). NR was supported by the Dutch Foundation for Applied Water Research (STOWA) and the waterboards Hoogheemraadschap Hollands Noorderkwartier and Waterschap De Dommel. KO was supported by the joint research program NWO–FAPESP of the Dutch Organization for Scientific Research (NWO) and the Sao Paulo Research Foundation (FAPESP) (code NWO: BBE.2017.013; code FAPESP: 2017/50249-6); and by the SIAM Gravitation Grant (024.002.002) from the Dutch Ministry of Education, Culture and Science (OCW).

## Notes

### Competing Interest Statement

The authors have declared no competing interest.

## References

1. Edenhofer O, Pichs-Madruga R, Sokona Y, Minx C, Farahani E, Kadner S, et al. Working Group III Contribution to the Fifth Assessment Report of the Intergovernmental Panel on Climate Change. Intergovernmental Panel on Climate Change: Cambridge, UK, New York, NY, USA. 2014.

2. Tian H, Xu R, Canadell JG, Thompson RL, Winiwarter W, Suntharalingam P, et al. A comprehensive quantification of global nitrous oxide sources and sinks. Nature. 2020;586(7828):248–56.

3. Thomson AJ, Giannopoulos G, Pretty J, Baggs EM, Richardson DJ. Biological sources and sinks of nitrous oxide and strategies to mitigate emissions. Philosophical Transactions of the Royal Society B: Biological Sciences. 2012;367(1593):1157–68.

4. Hallin S, Philippot L, Loffler FE, Sanford RA, Jones CM. Genomics and Ecology of Novel N2O-Reducing Microorganisms. Trends Microbiol. 2018;26(1):43–55.

5. Schreiber F, Wunderlin P, Udert KM, Wells GF. Nitric oxide and nitrous oxide turnover in natural and engineered microbial communities: biological pathways, chemical reactions, and novel technologies. Frontiers in Microbiology. 2012;3:372.

6. Wunderlin P, Mohn J, Joss A, Emmenegger L, Siegrist H. Mechanisms of N2O production in biological wastewater treatment under nitrifying and denitrifying conditions. Water Res. 2012;46(4):1027–37.

7. Simon J, Klotz MG. Diversity and evolution of bioenergetic systems involved in microbial nitrogen compound transformations. Biochim Biophys Acta. 2013;1827(2):114–35.

8. Domingo-Felez C, Smets BF. Modeling Denitrification as an Electric Circuit Accurately Captures Electron Competition between Individual Reductive Steps: The Activated Sludge Model-Electron Competition Model. Environ Sci Technol. 2020;54(12):7330–8.

9. Graf DR, Jones CM, Hallin S. Intergenomic comparisons highlight modularity of the denitrification pathway and underpin the importance of community structure for N2O emissions. PloS one. 2014;9(12):e114118.

10. Jones CM, Spor A, Brennan FP, Breuil M-C, Bru D, Lemanceau P, et al. Recently identified microbial guild mediates soil N2O sink capacity. Nature Climate Change. 2014;4(9):801–5.

11. Conthe M, Lycus P, Arntzen MO, Ramos da Silva A, Frostegard A, Bakken LR, et al. Denitrification as an N2O sink. Water Res. 2019;151:381–7.

12. Singleton CM, Petriglieri F, Kristensen JM, Kirkegaard RH, Michaelsen TY, Andersen MH, et al. Connecting structure to function with the recovery of over 1000 high-quality metagenome-assembled genomes from activated sludge using long-read sequencing. Nat Commun. 2021;12(1):2009.

13. Sanford RA, Wagner DD, Wu Q, Chee-Sanford JC, Thomas SH, Cruz-Garcia C, et al. Unexpected nondenitrifier nitrous oxide reductase gene diversity and abundance in soils. Proc Natl Acad Sci U S A. 2012;109(48):19709–14.

14. Jones CM, Graf DR, Bru D, Philippot L, Hallin S. The unaccounted yet abundant nitrous oxide-reducing microbial community: a potential nitrous oxide sink. ISME J. 2013;7(2):417–26.

15. Suenaga T, Hori T, Riya S, Hosomi M, Smets BF, Terada A. Enrichment, Isolation, and Characterization of High-Affinity N2O-Reducing Bacteria in a Gas-Permeable Membrane Reactor. Environ Sci Technol. 2019.

16. Yoon S, Nissen S, Park D, Sanford RA, Loffler FE. Nitrous Oxide Reduction Kinetics Distinguish Bacteria Harboring Clade I NosZ from Those Harboring Clade II NosZ. Appl Environ Microbiol. 2016;82(13):3793–800.

17. Marchant HK, Tegetmeyer HE, Ahmerkamp S, Holtappels M, Lavik G, Graf J, et al. Metabolic specialization of denitrifiers in permeable sediments controls N2 O emissions. Environ Microbiol. 2018;20(12):4486–502.

18. Bertagnolli AD, Konstantinidis KT, Stewart FJ. Non-denitrifier nitrous oxide reductases dominate marine biomes. Environ Microbiol Rep. 2020;12(6):681–92.

19. Herold M, Martinez Arbas S, Narayanasamy S, Sheik AR, Kleine-Borgmann LAK, Lebrun LA, et al. Integration of time-series meta-omics data reveals how microbial ecosystems respond to disturbance. Nat Commun. 2020;11(1):5281.

20. Lawson CE, Wu S, Bhattacharjee AS, Hamilton JJ, McMahon KD, Goel R, et al. Metabolic network analysis reveals microbial community interactions in anammox granules. Nat Commun. 2017;8:15416.

21. Speth DR, I. ‘t Zandt MH, Guerrero-Cruz S, Dutilh BE, Jetten MS. Genome-based microbial ecology of anammox granules in a full-scale wastewater treatment system. Nat Commun. 2016;7:11172.

22. Torresi E, Gulay A, Polesel F, Jensen MM, Christensson M, Smets BF, et al. Reactor staging influences microbial community composition and diversity of denitrifying MBBRs-Implications on pharmaceutical removal. Water Res. 2018;138:333–45.

23. Vieira A, Galinha CF, Oehmen A, Carvalho G. The link between nitrous oxide emissions, microbial community profile and function from three full-scale WWTPs. The Science of the total environment. 2019;651(Pt 2):2460–72.

24. Domingo-Félez C, Smets BF. A consilience model to describe N2O production during biological N removal. Environmental Science: Water Research & Technology. 2016;2(6):923–30.

25. Mania D, Woliy K, Degefu T, Frostegard A. A common mechanism for efficient N(2) O reduction in diverse isolates of nodule-forming bradyrhizobia. Environ Microbiol. 2020;22(1):17–31.

26. Yoon H, Song MJ, Kim DD, Sabba F, Yoon S. A Serial Biofiltration System for Effective Removal of Low-Concentration Nitrous Oxide in Oxic Gas Streams: Mathematical Modeling of Reactor Performance and Experimental Validation. Environ Sci Technol. 2019;53(4):2063–74.

27. Yoon H, Song MJ, Yoon S. Design and Feasibility Analysis of a Self-Sustaining Biofiltration System for Removal of Low Concentration N2O Emitted from Wastewater Treatment Plants. Environ Sci Technol. 2017;51(18):10736–45.

28. Conthe M, Wittorf L, Kuenen JG, Kleerebezem R, Hallin S, van Loosdrecht MCM. Growth yield and selection of nosZ clade II types in a continuous enrichment culture of N2O respiring bacteria. Environ Microbiol Rep. 2018;10(3):239–44.

29. Conthe M, Wittorf L, Kuenen JG, Kleerebezem R, van Loosdrecht MCM, Hallin S. Life on N2O: deciphering the ecophysiology of N2O respiring bacterial communities in a continuous culture. ISME J. 2018;12(4):1142–53.

30. Qi C, Zhou Y, Suenaga T, Oba K, Lu J, Wang G, et al. Organic carbon determines nitrous oxide consumption activity of clade I and II nosZ bacteria: Genomic and biokinetic insights. Water Res. 2021;209:117910.

31. Semedo M, Wittorf L, Hallin S, Song B. Differential expression of clade I and II N2O reductase genes in denitrifying Thauera linaloolentis 47LolT under different nitrogen conditions. FEMS Microbiol Lett. 2020;367(24).

32. LaSarre B, Morlen R, Neumann GC, Harwood CS, McKinlay JB. Nitrous oxide reduction by two partial denitrifying bacteria requires denitrification intermediates that cannot be respired. 2024;90(1):e01741–23.

33. van der Star WR, van de Graaf MJ, Kartal B, Picioreanu C, Jetten MS, van Loosdrecht MC. Response of anaerobic ammonium-oxidizing bacteria to hydroxylamine. Appl Environ Microbiol. 2008;74(14):4417–26.

34. Vishniac W, Santer MJBr. The thiobacilli. 1957;21(3):195–213.

35. APHA. Standard methods for the examination of water and wastewater. Washington, D.C. 2005.

36. Roels J. Energetics and kinetics in biotechnology: Elsevier Biomedical Press; 1983.

37. Hellinga C, Romein B. MACROBAL: A Program for Robust Data Reconciliation and Gross Error Detection. IFAC Proceedings Volumes. 1992;25(2):459–60.

38. Sander R. Compilation of Henry’s law constants (version 4.0) for water as solvent. Atmospheric Chemistry and Physics. 2015;15(8):4399–981.

39. Bolger AM, Lohse M, Usadel B. Trimmomatic: a flexible trimmer for Illumina sequence data. Bioinformatics. 2014;30(15):2114–20.

40. Nurk S, Meleshko D, Korobeynikov A, Pevzner PA. metaSPAdes: a new versatile metagenomic assembler. Genome Res. 2017;27(5):824–34.

41. Li H. Aligning sequence reads, clone sequences and assembly contigs with BWA-MEM. arXiv preprint arXiv:13033997. 2013.

42. Li H, Handsaker B, Wysoker A, Fennell T, Ruan J, Homer N, et al. The sequence alignment/map format and SAMtools. Bioinformatics. 2009;25(16):2078–9.

43. Kang DD, Li F, Kirton E, Thomas A, Egan R, An H, et al. MetaBAT 2: an adaptive binning algorithm for robust and efficient genome reconstruction from metagenome assemblies. PeerJ. 2019;7:e7359.

44. Parks DH, Imelfort M, Skennerton CT, Hugenholtz P, Tyson GW. CheckM: assessing the quality of microbial genomes recovered from isolates, single cells, and metagenomes. Genome research. 2015;25(7):1043–55.

45. Parks DH, Rinke C, Chuvochina M, Chaumeil PA, Woodcroft BJ, Evans PN, et al. Recovery of nearly 8,000 metagenome-assembled genomes substantially expands the tree of life. Nat Microbiol. 2017;2(11):1533–42.

46. Chaumeil PA, Mussig AJ, Hugenholtz P, Parks DH. GTDB-Tk v2: memory friendly classification with the genome taxonomy database. Bioinformatics. 2022;38(23):5315–6.

47. Dong X, Strous M. An Integrated Pipeline for Annotation and Visualization of Metagenomic Contigs. Frontiers in Genetics. 2019;10.

48. Kanehisa M. Enzyme Annotation and Metabolic Reconstruction Using KEGG. Methods Mol Biol. 2017;1611:135–45.

49. Paysan-Lafosse T, Blum M, Chuguransky S, Grego T, Pinto BL, Salazar GA, et al. InterPro in 2022. Nucleic Acids Res. 2023;51(D1):D418–D27.

50. Simon J, Einsle O, Kroneck PM, Zumft WG. The unprecedented nos gene cluster of Wolinella succinogenes encodes a novel respiratory electron transfer pathway to cytochrome c nitrous oxide reductase. FEBS Lett. 2004;569(1-3):7–12.

51. Zallot R, Ross R, Chen WH, Bruner SD, Limbach PA, de Crecy-Lagard V. Identification of a Novel Epoxyqueuosine Reductase Family by Comparative Genomics. ACS Chem Biol. 2017;12(3):844–51.

52. Apweiler R, Bairoch A, Wu CH, Barker WC, Boeckmann B, Ferro S, et al. UniProt: the Universal Protein knowledgebase. Nucleic Acids Res. 2004;32(Database issue):D115–9.

53. RStudio Team. RStudio: Integrated Development Environment for R. RStudio, PBC, 777 Boston, MA http://www.rstudio.com/. 2021.

54. R Core Team. R: A language and environment for statistical computing. R Foundation 779 for Statistical Computing, Vienna, Austria https://www.r-project.org/. 2022.

55. Banks RGS, Henderson RJ, Pratt JM. Reactions of gases in solution. Part III. Some reactions of nitrous oxide with transition-metal complexes. Journal of the Chemical Society A: Inorganic, Physical, Theoretical. 1968(0):2886–9.

56. Deacon R, Perry J, Lumb M, Chanarin I, Minty B, Halsey MJ, et al. Selective inactivation of vitamin B12 in rats by nitrous oxide. The Lancet. 1978;312(8098):1023–4.

57. Sullivan MJ, Gates AJ, Appia-Ayme C, Rowley G, Richardson DJ. Copper control of bacterial nitrous oxide emission and its impact on vitamin B12-dependent metabolism. Proc Natl Acad Sci U S A. 2013;110(49):19926–31.

58. Kleiner M, Thorson E, Sharp CE, Dong X, Liu D, Li C, et al. Assessing species biomass contributions in microbial communities via metaproteomics. Nat Commun. 2017;8(1):1558.

59. Barco RA, Garrity GM, Scott JJ, Amend JP, Nealson KH, Emerson D. A Genus Definition for Bacteria and Archaea Based on a Standard Genome Relatedness Index. mBio. 2020;11(1):10.1128/mbio.02475-19.

60. Thomsen TR, Kong Y, Nielsen PH. Ecophysiology of abundant denitrifying bacteria in activated sludge. FEMS Microbiol Ecol. 2007;60(3):370–82.

61. Wang Z, Li W, Li H, Zheng W, Guo F. Phylogenomics of Rhodocyclales and its distribution in wastewater treatment systems. Scientific reports. 2020;10(1):3883.

62. Wu L, Ning D, Zhang B, Li Y, Zhang P, Shan X, et al. Global diversity and biogeography of bacterial communities in wastewater treatment plants. Nat Microbiol. 2019;4(7):1183–95.

63. Conthe M, Parchen C, Stouten G, Kleerebezem R, van Loosdrecht MCM. O2 versus N2O respiration in a continuous microbial enrichment. Appl Microbiol Biotechnol. 2018;102(20):8943–50.

64. Kim DD, Han H, Yun T, Song MJ, Terada A, Laureni M, et al. Identification of nosZ-expressing microorganisms consuming trace N2O in microaerobic chemostat consortia dominated by an uncultured Burkholderiales. ISME J. 2022.

65. Jonassen KR. Biogas digestate as substrate and vector for the introduction of N2O-respiring bacteria to agricultural soil.: Norwegian University of Life Sciences; 2021.

66. Valk LC, Peces M, Singleton CM, Laursen MD, Andersen MH, Mielczarek AT, et al. Exploring the microbial influence on seasonal nitrous oxide concentration in a full-scale wastewater treatment plant using metagenome assembled genomes. Water Res. 2022;219:118563.

67. van den Berg EM, van Dongen U, Abbas B, van Loosdrecht MC. Enrichment of DNRA bacteria in a continuous culture. ISME J. 2015;9(10):2153–61.

68. Zumft WG. Chapter 13 - Respiratory Nitric Oxide Reductases, NorB and NorZ, of the Heme– Copper Oxidase Type. In: Ghosh A, editor. The Smallest Biomolecules: Diatomics and their Interactions with Heme Proteins. Amsterdam: Elsevier; 2008. p. 327–53.

69. Roth JR, Lawrence JG, Bobik TA. Cobalamin (coenzyme B12): synthesis and biological significance. Annu Rev Microbiol. 1996;50:137–81.

70. Lu X, Heal KR, Ingalls AE, Doxey AC, Neufeld JD. Metagenomic and chemical characterization of soil cobalamin production. ISME J. 2020;14(1):53–66.

71. Shelton AN, Seth EC, Mok KC, Han AW, Jackson SN, Haft DR, et al. Uneven distribution of cobamide biosynthesis and dependence in bacteria predicted by comparative genomics. ISME J. 2019;13(3):789–804.

72. Lu J, Tappel RC, Nomura CT. Mini-Review: Biosynthesis of Poly(hydroxyalkanoates). Polymer Reviews. 2009;49(3):226–48.

73. Morris JJ, Lenski RE, Zinser ER. The Black Queen Hypothesis: evolution of dependencies through adaptive gene loss. mBio. 2012;3(2).

74. Bjorkeroth J, Campbell K, Malina C, Yu R, Di Bartolomeo F, Nielsen J. Proteome reallocation from amino acid biosynthesis to ribosomes enables yeast to grow faster in rich media. Proc Natl Acad Sci U S A. 2020;117(35):21804–12.

75. O’Brien EJ, Utrilla J, Palsson BO. Quantification and Classification of E. coli Proteome Utilization and Unused Protein Costs across Environments. PLoS Comput Biol. 2016;12(6):e1004998.

76. Peebo K, Valgepea K, Maser A, Nahku R, Adamberg K, Vilu R. Proteome reallocation in Escherichia coli with increasing specific growth rate. Mol Biosyst. 2015;11(4):1184–93.

77. Men Y, Yu K, Baelum J, Gao Y, Tremblay J, Prestat E, et al. Metagenomic and Metatranscriptomic Analyses Reveal the Structure and Dynamics of a Dechlorinating Community Containing Dehalococcoides mccartyi and Corrinoid-Providing Microorganisms under Cobalamin-Limited Conditions. Appl Environ Microbiol. 2017;83(8).

