## Supplementary Information for "Selective enrichment of high-affinity clade II N_2_O-reducers in a mixed culture"

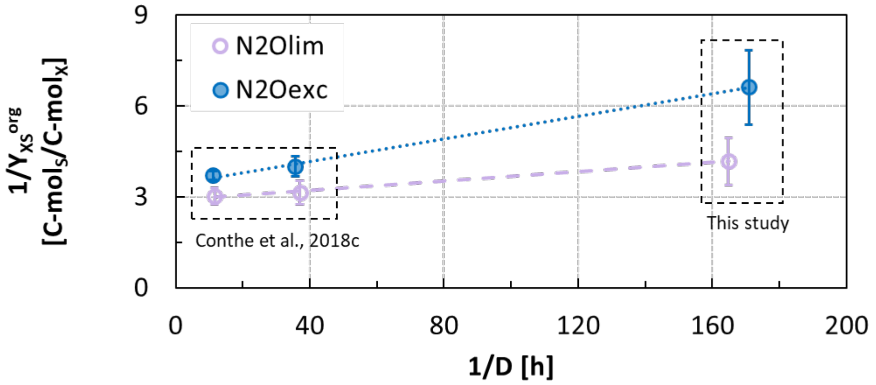


**Figure S1**. Estimation of the biomass-specific acetate consumption rate for maintenance based on the estimated growth yields on acetate over the dilution rates range covered by this study and (1).

**Table S1**. Average and standard deviation of the dilution rate (D), and the conversion rates for the two enrichments. Values are calculated from the steady-state reactor operation measurements (Figure 1; days 100-150 for *$N_2Oexc$* and 60-100 for *$N_2Olim$*). The software Macrobal (2) was used for data reconciliation.

| **D** | **Acetate** | **N2O** | **NH4+** | **X** | **H+** | |
|---|---|---|---|---|---|---|
| **$h^{-1}$** | **mmol/d** | **mmol/d** | **mmol/d** | **mmol/d** | **mmol/d** | |
| 0.0059 ± 0.001 | -30.6 ± 1.8 | -104.7 ± 8.8 | -3.8 ± 1.2 | 9.3 ± 1.6 | -26.8 ± 2 | **$N_2Oexc$** |
| 0.0061 ± 0.0007 | -24.4 ± 1.1 | -75.6 ± 1.6 | -5.1 ± 1.2 | 11.7 ± 2.1 | -19.2 ± 1.5 | **$N_2Olim$** |

**Table S2.** Full taxonomy of the recovered bins.

| | Bin ID | Phylum | Class | Order | Family | Genus | Species |
|---|---|---|---|---|---|---|---|
| $N_2Oexc$ | **Bin.820_exc** | Pseudomonadota | Gammaproteobacteria | Burkholderiales | Rhodocyclaceae | Azonexus | |
| | **Bin.2_exc** | Pseudomonadota | Alphaproteobacteria | Rhodobacterales | Rhodobacteraceae | Phaeovulum | |
| | **Bin.24_exc** | Pseudomonadota | Alphaproteobacteria | Rhodobacterales | Rhodobacteraceae | Pseudorhodobacter | |
| | **Bin.15_exc** | Bdellovibrionota | Bdellovibrionia | Bdellovibrionales | Bdellovibrionaceae | Bdellovibrio | Bdellovibrio sp019104905 |
| | **Bin.3_exc** | Pseudomonadota | Alphaproteobacteria | Rhizobiales | Rhizobiaceae | Aquamicrobium | |
| | **Bin.12_exc** | Bacteroidota | Bacteroidia | Flavobacteriales | Flavobacteriaceae | Flavobacterium | Flavobacterium filum |
| | **Bin.18_exc** | Bacteroidota | Bacteroidia | Flavobacteriales | Weeksellaceae | Moheibacter | Moheibacter sp019455045 |
| | **Bin.16_exc** | Bacteroidota | Bacteroidia | Flavobacteriales | Weeksellaceae | Kaistella | |
| | **Bin.17_exc** | Bacteroidota | Bacteroidia | Chitinophagales | Chitinophagaceae | Ferruginibacter | Ferruginibacter sp001898465 |
| | **Bin.9_exc** | Bacteroidota | Bacteroidia | Chitinophagales | Chitinophagaceae | Ferruginibacter | Ferruginibacter sp018268175 |
| | ***unbinned*** | | | | | | |
| $N_2Olim$ | **Bin.9_lim** | Pseudomonadota | Gammaproteobacteria | Burkholderiales | Rhodocyclaceae | Azonexus | Azonexus sp900549295 |
| | **Bin.16_lim** | Pseudomonadota | Gammaproteobacteria | Burkholderiales | Rhodocyclaceae | Thauera | Thauera phenylacetica |
| | **Bin.14_lim** | Bacteroidota | Bacteroidia | Flavobacteriales | Weeksellaceae | UBA3376 | |
| | **Bin.21_lim** | Bacteroidota | Bacteroidia | Chitinophagales | Chitinophagaceae | Ferruginibacter | Ferruginibacter sp018268175 |
| | **Bin.6_lim** | Deferribacterota | Deferribacteres | Deferribacterales | Denitrovibrionaceae | Seleniivibrio | Seleniivibrio woodruffii |
| | **Bin.24_lim** | Patescibacteria | JAEDAM01 | BD1-5 | UBA6164 | UBA7396 | |
| | **Bin.20._lim** | Pseudomonadota | Gammaproteobacteria | Burkholderiales | Rhodocyclaceae | Azovibrio | Azovibrio restrictus |
| | **Bin.26_lim** | Bacillota | Clostridia | Oscillospirales | Oscillospiraceae | Pseudoflavonifractor | |
| | **Bin.11_lim** | Bacteroidota | Bacteroidia | Bacteroidales | F082 | JALNZU01 | JALNZU01 sp002421995 |
| | **Bin.2_lim** | Bacillota | Clostridia | Peptostreptococcales | Anaerovoracaceae | UBA5559 | |
| | **Bin.8_lim** | Bacteroidota | Bacteroidia | Chitinophagales | Chitinophagaceae | Niabella | |
| | **Bin.10_lim** | Bacteroidota | Bacteroidia | Bacteroidales | WCHB1-69 | UBA5429 | |
| | **Bin.1_lim** | Bacillota | Bacilli | Acholeplasmatales | UBA5453 | Paracholeplasma | |
| | **Bin.27_lim** | Bacillota | Clostridia | Peptostreptococcales | Acidaminobacteraceae | Fusibacter_C | Fusibacter_C sp001897605 |
| | **Bin.23_lim** | Bacteroidota | Bacteroidia | Bacteroidales | P3 | UBA10566 | UBA10566 sp002399565 |
| | **Bin.22_lim** | Bacillota | Bacilli | Erysipelotrichales | Erysipelotrichaceae | UBA2212 | |
| | **Bin.18_lim** | Bacteroidota | Bacteroidia | Bacteroidales | Paludibacteraceae | UPXZ01 | |
| | ***unbinned*** | | | | | | |

**Table S3**. Annotation of the enzymes involved in cobalamin synthesis and transport, and the cobalamin-dependent enzymes with their cobalamin-independent functional homologues.

| ko | ko_name | EC_1 | EC_2 | EC_3 | name | pathway |
|---|---|---|---|---|---|---|
| K02492 | hemA | 1.2.1.70 | | | hemA | Siroheme biosynthesis |
| K01845 | hemL | 5.4.3.8 | | | hemL | Siroheme biosynthesis |
| K01698 | hemB, ALAD | 4.2.1.24 | | | hemB, ALAD | Siroheme biosynthesis |
| K01749 | hemC, HMBS | 2.5.1.61 | | | hemC, HMBS | Siroheme biosynthesis |
| K01719 | hemD, UROS | 4.2.1.75 | | | hemD, UROS | Siroheme biosynthesis |
| K02302 | cysG | 2.1.1.107 | 1.3.1.76 | 4.99.1.4 | hemDX, hemX, MET1, cobA, cobA-hemD, cysG | Siroheme biosynthesis |
| K02303 | cobA | 2.1.1.107 | | | hemDX, hemX, MET1, cobA, cobA-hemD, cysG | Siroheme biosynthesis |
| K13542 | cobA-hemD | 2.1.1.107 | 4.2.1.75 | | hemDX, hemX, MET1, cobA, cobA-hemD, cysG | Siroheme biosynthesis |
| K00589 | MET1 | 2.1.1.107 | | | hemDX, hemX, MET1, cobA, cobA-hemD, cysG | Siroheme biosynthesis |
| K02496 | hemX | 2.1.1.107 | | | hemDX, hemX, MET1, cobA, cobA-hemD, cysG | Siroheme biosynthesis |
| K13543 | hemDX | 2.1.1.107 | 4.2.1.75 | | hemDX, hemX, MET1, cobA, cobA-hemD, cysG | Siroheme biosynthesis |
| K02304 | MET8 | 1.3.1.76 | 4.99.1.4 | | MET8, sirC | Siroheme biosynthesis |
| K24866 | sirC | 1.3.1.76 | | | MET8, sirC | Siroheme biosynthesis |
| K03794 | sirB | 4.99.1.4 | | | sirB | Siroheme biosynthesis |
| K02190 | cbiK | 4.99.1.3 | | | cbiK, cbiX, cfbA | Corrin ring biosynthesis (aerobic/aerobic) |
| K03795 | cbiX | 4.99.1.3 | | | cbiK, cbiX, cfbA | Corrin ring biosynthesis (aerobic/aerobic) |
| K22011 | cfbA | 4.99.1.3 | 4.99.1.11 | | cbiK, cbiX, cfbA | Corrin ring biosynthesis (aerobic/aerobic) |
| K03394 | cobI-cbiL | 2.1.1.130 | 2.1.1.151 | | cobI-cbiL, cobIJ | Corrin ring biosynthesis (aerobic/aerobic) |
| K13540 | cobIJ | 2.1.1.130 | 2.1.1.131 | | cobI-cbiL, cobIJ | Corrin ring biosynthesis (aerobic/aerobic) |
| K02229 | cobG | 1.14.13.83 | | | cobG | Corrin ring biosynthesis (aerobic/aerobic) |
| K05934 | cobJ, cbiH | 2.1.1.272 | 2.1.1.131 | | cbiH60, cbiGH-cobJ, cobJ, cbiH | Corrin ring biosynthesis (aerobic/aerobic) |
| K13541 | cbiGH-cobJ | 2.1.1.272 | 2.1.1.131 | 3.7.1.12 | cbiH60, cbiGH-cobJ, cobJ, cbiH | Corrin ring biosynthesis (aerobic/aerobic) |
| K21479 | cbiH60 | 2.1.1.272 | | | cbiH60, cbiGH-cobJ, cobJ, cbiH | Corrin ring biosynthesis (aerobic/aerobic) |
| K05936 | cobM, cbiF | 2.1.1.133 | 2.1.1.271 | | cobM, cbiF | Corrin ring biosynthesis (aerobic/aerobic) |
| K02189 | cbiG | 3.7.1.12 | | | cbiG | Corrin ring biosynthesis (aerobic/aerobic) |
| K02188 | cbiD | 2.1.1.195 | | | cbiD | Corrin ring biosynthesis (aerobic/aerobic) |
| K02228 | cobF | 2.1.1.152 | | | cobF | Corrin ring biosynthesis (aerobic/aerobic) |
| K05895 | cobK-cbiJ | 1.3.1.54 | 1.3.1.106 | | cobK-cbiJ | Corrin ring biosynthesis (aerobic/aerobic) |
| K02191 | cbiT | 2.1.1.196 | | | cbiT | Corrin ring biosynthesis (aerobic/aerobic) |
| K00595 | cobL-cbiET | 2.1.1.289 | 2.1.1.196 | 2.1.1.132 | cobL-cbiET, cbiE | Corrin ring biosynthesis (aerobic/aerobic) |
| K03399 | cbiE | 2.1.1.289 | | | cobL-cbiET, cbiE | Corrin ring biosynthesis (aerobic/aerobic) |
| K06042 | cobH-cbiC | 5.4.99.61 | 5.4.99.60 | | cobH-cbiC | Corrin ring biosynthesis (aerobic/aerobic) |
| K02224 | cobB-cbiA | 6.3.5.9 | 6.3.5.11 | | cobB-cbiA | Corrin ring biosynthesis (aerobic/aerobic) |
| K02230 | cobN | 6.6.1.2 | | | cobN, cobS, cobT | Corrin ring biosynthesis (aerobic/aerobic) |
| K09882 | cobS | 6.6.1.2 | | | cobN, cobS, cobT | Corrin ring biosynthesis (aerobic/aerobic) |

| | | | | | |
|---|---|---|---|---|---|
| K09883 | cobT | 6.6.1.2 | | cobN, cobS, cobT | Corrin ring biosynthesis (aerobic/aerobic) |
| K13786 | cobR | | | cobR | Nucleotide loop assembly |
| K00798 | MMAB, pduO | 2.5.1.17 | | MMAB, pduO, cobA, btuR | Nucleotide loop assembly |
| K19221 | cobA, btuR | 2.5.1.17 | | MMAB, pduO, cobA, btuR | Nucleotide loop assembly |
| K02232 | cobQ, cbiP | 6.3.5.10 | | cobQ, cbiP | Nucleotide loop assembly |
| K02225 | cobC1, cobC | 6.3.1.10 | | cobC1, cobC, cbiB, cobD | Nucleotide loop assembly |
| K02227 | cbiB, cobD | 6.3.1.10 | | cobC1, cobC, cbiB, cobD | Nucleotide loop assembly |
| K02231 | cobP, cobU | 2.7.1.156 | | cobP, cobU | Nucleotide loop assembly |
| K19712 | cobY | 2.7.7.62 | | cobY | Nucleotide loop assembly |
| K00768 | cobU, cobT | 2.4.2.21 | | cobU, cobT | Nucleotide loop assembly |
| K02226 | cobC, phpB | 3.1.3.73 | | rh-cobC, cobC, phpB | Nucleotide loop assembly |
| K22316 | rh-cobC | 3.1.3.73 | 3.1.26.4 | rh-cobC, cobC, phpB | Nucleotide loop assembly |
| K02233 | cobS, cobV | 2.7.8.26 | | cobS, cobV | Nucleotide loop assembly |
| K16092 | btuB | | | btuB | B12 transporter |
| K00548 | metH, MTR | 2.1.1.13 | | (B12-dependent) yitJ, metH | Methionine synthase |
| K24042 | yitJ | 2.1.1.13 | 1.5.1.54 | (B12-dependent) yitJ, metH | Methionine synthase |
| K00549 | metE | 2.1.1.14 | | (B12-independent) metE | Methionine synthase |
| K18979 | queG | 1.17.99.6 | | (B12-dependent) queG | Epoxyqueuosine reductase |
| K09765 | queH | 1.17.99.6 | | (B12-independent) queH | Epoxyqueuosine reductase |
| K00524 | nrdJ | 1.17.4.2 | | (B12-dependent) rtpR, nrdJ | Ribonucleoside-triphosphate reductase |
| K00527 | rtpR | 1.17.4.2 | | (B12-dependent) rtpR, nrdJ | Ribonucleoside-triphosphate reductase |
| K21636 | nrdD | 1.1.98.6 | | (B12-independent) nrdD | Ribonucleoside-triphosphate reductase |
| K01847 | MUT | 5.4.99.2 | | (B12-dependent) MUT | Methylmalonyl-CoA mutase |

### References


1. Conthe M, Wittorf L, Kuenen JG, Kleerebezem R, van Loosdrecht MCM, Hallin S. Life on N2O: deciphering the ecophysiology of N2O respiring bacterial communities in a continuous culture. ISME J. 2018;12(4):1142-53.
2. Hellinga C, Romein B. MACROBAL: A Program for Robust Data Reconciliation and Gross Error Detection. IFAC Proceedings Volumes. 1992;25(2):459-60.
